# SARITA: A Large Language Model for Generating the S1 Subunit of the SARS-CoV-2 Spike Protein

**DOI:** 10.1101/2024.12.10.627777

**Authors:** Simone Rancati, Giovanna Nicora, Laura Bergomi, Tommaso Mario Buonocore, Daniel M Czyz, Enea Parimbelli, Riccardo Bellazzi, Marco Salemi, Mattia Prosperi, Simone Marini

## Abstract

The COVID-19 pandemic has profoundly impacted global health, economics, and daily life, with over 776 million cases and 7 million deaths from December 2019 to November 2024. Since the original SARS-CoV-2 Wuhan strain emerged, the virus has evolved into variants such as Alpha, Beta, Gamma, Delta, and Omicron, all characterized by mutations in the Spike glycoprotein, critical for viral entry into human cells via its S1 and S2 subunits. The S1 subunit, binding to the ACE2 receptor and mutating frequently, affects infectivity and immune evasion; the more conserved S2, on the other hand, facilitates membrane fusion. Predicting future mutations is crucial for developing vaccines and treatments adaptable to emerging strains, enhancing preparedness and intervention design. Generative Large Language Models (LLMs) are becoming increasingly common in the field of genomics, given their ability to generate realistic synthetic biological sequences, including applications in protein design and engineering. Here we present SARITA, an LLM with up to 1.2 billion parameters, based on GPT-3 architecture, designed to generate high-quality synthetic SARS-CoV-2 Spike S1 sequences. SARITA is trained via continuous learning on the pre-existing protein model RITA. When trained on Alpha, Beta, and Gamma variants (data up to February 2021 included), SARITA correctly predicts the evolution of future S1 mutations, including characterized mutations of Delta, Omicron and Iota variants. Furthermore, we show how SARITA outperforms alternative approaches, including other LLMs, in terms of sequence quality, realism, and similarity with real-world S1 sequences. These results indicate the potential of SARITA to predict future SARS-CoV-2 S1 evolution, potentially aiding in the development of adaptable vaccines and treatments.

## Introduction

The COVID-19 pandemic, caused by the SARS-CoV-2, has significantly affected global health, economies, and daily human life, resulting in over 776 million estimated cases and 7 million deaths between December 2019 and November 2024 [1]. Since the emergence of the original Wuhan strain, SARS-CoV-2 has evolved into variants such as Alpha, Beta, Gamma, Delta, and Omicron, all distinguished by mutations in the Spike glycoprotein (**Figure 1A**). The Spike protein is vital for the virus entry into human cells via its S1 and S2 subunits (**Figure 1B**) [2]. More specifically, the S1 subunit (672 amino acids long) is more mutable and contains the receptor-binding domain (RBD), which recognizes and binds to the host angiotensin-converting enzyme 2 (ACE2). This subunit is crucial for the initial binding of the virus to host cells and shows significant variability, making it a key target for neutralizing antibodies. However, S1 variability can also hinder the development of broad-spectrum antiviral drugs, as mutations may alter its conformation and reduce the efficacy of antibodies. The S2 subunit (587 amino acids long), on the other hand, is more conserved and mediates the fusion of the viral membrane with that of the host cell through the formation of a six-helix bundle. This region is less prone to mutations compared to S1, and thus provides a more stable target for the development of fusion inhibitors [1], [2].

**Figure 1:**
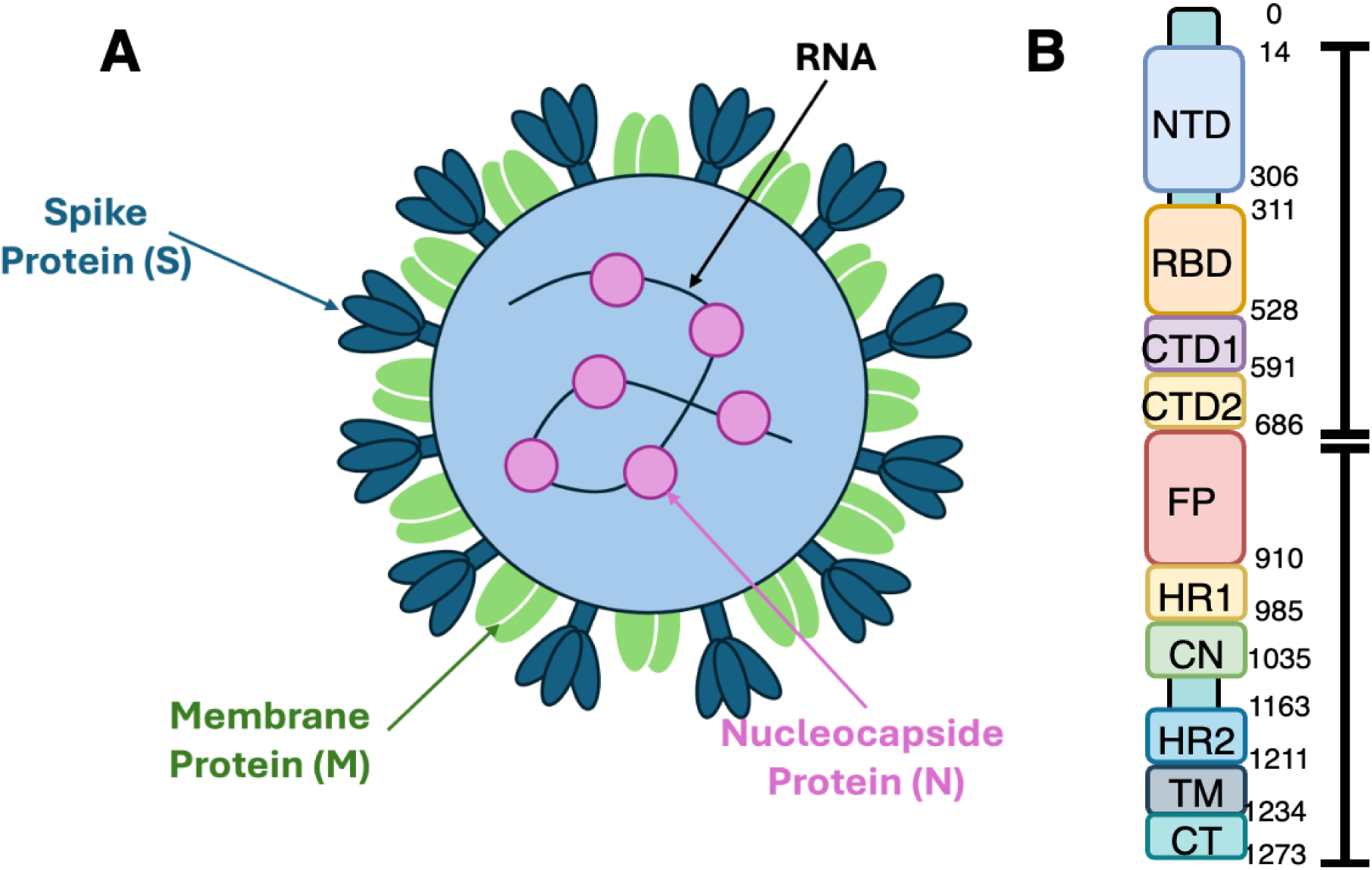
A) **Schematic representation of the SARS-CoV-2 virus**, showing major structural components relevant for virus replication and entry into the host cell. The RNA genome (pink) is tightly associated with the nucleocapsid protein (N, purple) and is wrapped inside the virus. These structural components are functional for the replication cycle of the virus and its structural integrity. The membrane protein (M, green) provides for the virus shape preservation and the assembly of new viral particles. Projecting from the viral surface are Spike proteins (S, blue), essential for binding to host cell receptors and facilitating cell entry; as such, Spike proteins represent a key target for vaccine and therapeutic development. B) **Schematic Representation of SARS-CoV-2 Spike Protein:** The diagram illustrates the Spike protein structure divided into the S1 and S2 subunits, each containing specific domains. The S1 subunit includes the N-terminal domain (NTD), the receptor-binding domain (RBD), and two C-terminal domains (CTD1 and CTD2). In the S2 subunit, highly conserved, notable features include the S2’ cleavage site, the fusion peptide (FP), heptad repeat regions (HR1 and HR2), a central helix region (CH), and a connector domain (CD). The S1/S2 cleavage between CTD2 and FP indicates the enzymatic division point between the subunits. The diagram also shows the transmembrane anchor (TM) and the cytoplasmic tail (CT), which anchor the protein to the virus membrane. Glycans, depicted as tree-like symbols, are attached to specified sites, playing roles in protein folding and immune evasion. (Image realized with draw.io [3].)

In the SARS-CoV-2 ever-evolving genomic landscape, accurately predicting viral evolution is essential for implementing effective public health measures and developing new vaccines and therapies. Various methods have been devised to forecast whether a current SARS-CoV-2 sequence might become a variant of concern [4], [5] or a dominant lineage in the future [6], or to predict the likelihood of a given viral protein variant to induce immune escape from antibodies [7]. While these bioinformatics tools are critical to quickly respond to viral evolution, especially if used in the context of genomic surveillance, they can detect new, dangerous variants only post-sequencing, i.e., *after* the virus has already evolved them. In other words, these methods allow us only to react to viral evolution rather than proactively countering it. If we want to shift to a more proactive stance and anticipate the virus evolution, we must develop tools capable of predicting dangerous variants *before* they emerge, and preemptively design appropriate treatments and vaccines.

Methods based on Large Language Models (LLMs), may accurately capture the biological context of a protein sequence [8], [9], unlike simpler approaches like *k*-mer representations, which inherently overlook the positional information of (sub)sequences and amino acids, or substitution matrices such as PAM and BLOSUM [10] , which rely on statistical properties only. Recent research has investigated the use of LLMs to predict and generate novel SARS-CoV-2 sequences. Dhodapkar et al. introduced SpikeGPT2, a model fine-tuned on Spike protein sequences, which successfully forecasted future RBD (∼100 amino acids) mutations and identified potentially high-risk variants in this subsection of the Spike protein [11]. The main limit of SpikeGPT2 is that, while trained on the whole Spike protein, it is tested only on the RBD specific subsection. Ramachandran et al. proposed PandoGen, a method combining synthetic data generation, conditional sequence generation, and reward-based learning to forecast future SARS-CoV-2 sequences [12]. However, while the PandoGen code is accessible, the model is not readily available, as the user should reproduce the whole fine process, requiring both V100 and A100 GPUs (4 GPUs in total) in order to obtain the model. Additionally, these models have a large number of parameters (738 millions for SpikeGPT2; 192 millions for PandoGen), which makes them complex and difficult to use in settings where computing power is limited [11], [12].

To fulfill this research gap we develop SARITA (or SARS-CoV-2 RITA), an LLM designed to generate new, synthetic, high-quality and highly realistic SARS-CoV-2 S1 subunits. SARITA builds upon the continual learning framework of RITA, a state-of-the-art generative language model [13]. RITA is an autoregressive model for general protein sequence generation with up to 1.2 billion parameters. To capture the unique biological features of the Spike protein and obtain a specialized approach, we apply continual learning to pre-train RITA via 612,759 high-quality SARS-CoV-2 S1 sequences from GISAID [14]. To match different needs in terms of computational capacities, SARITA comes in four different sizes: The smallest model has 85 million parameters, while the largest has 1.2 billion. SARITA generates new S1 sequences using as an input the 14 amino acid sequence preceding it. We show how SARITA provides significantly better performances than concurrent models, including other LLMs, not only in generating high-quality and realistic S1 sequences, but also precisely predicting the emergence of 2022-2023 Spike mutations when trained on data from 2019 to 2021.

## Methods

### Architecture

The SARITA architecture (**Figure 2**) is based on a series of decoder-only transformers, inspired by the GPT-3 model [15]. It employs Rotary Positional Embeddings (RoPE) to enhance the model’s ability to capture positional relationships within the input data [16]. SARITA is available in four configurations: SARITA-S with 85 million parameters, featuring an embedding size of 768 and 12 transformer layers; SARITA-M with 300 million parameters, featuring an embedding dimension of 1024 and 24 layers; SARITA-L with 680 million parameters featuring an embedding size of 1536 and 24 layers; and SARITA-XL, with 1.2 billion parameters, featuring an embedding size of 2048, and 24 layers. All SARITA models can generate sequences up to 1024 tokens long. SARITA uses the Unigram model for tokenization, where each amino acid is represented as a single token, reflecting its unique role in protein structure and function. The tokenizer also includes special tokens like <PAD> for padding shorter sequences and <EOS> for marking sequence ends, ensuring consistency across datasets. This process reduces variability and enhances the model’s ability to learn meaningful patterns from protein sequences [13]. At the end each token is transformed into a numerical representation using a look-up table (Table S1).

**Figure 2:**
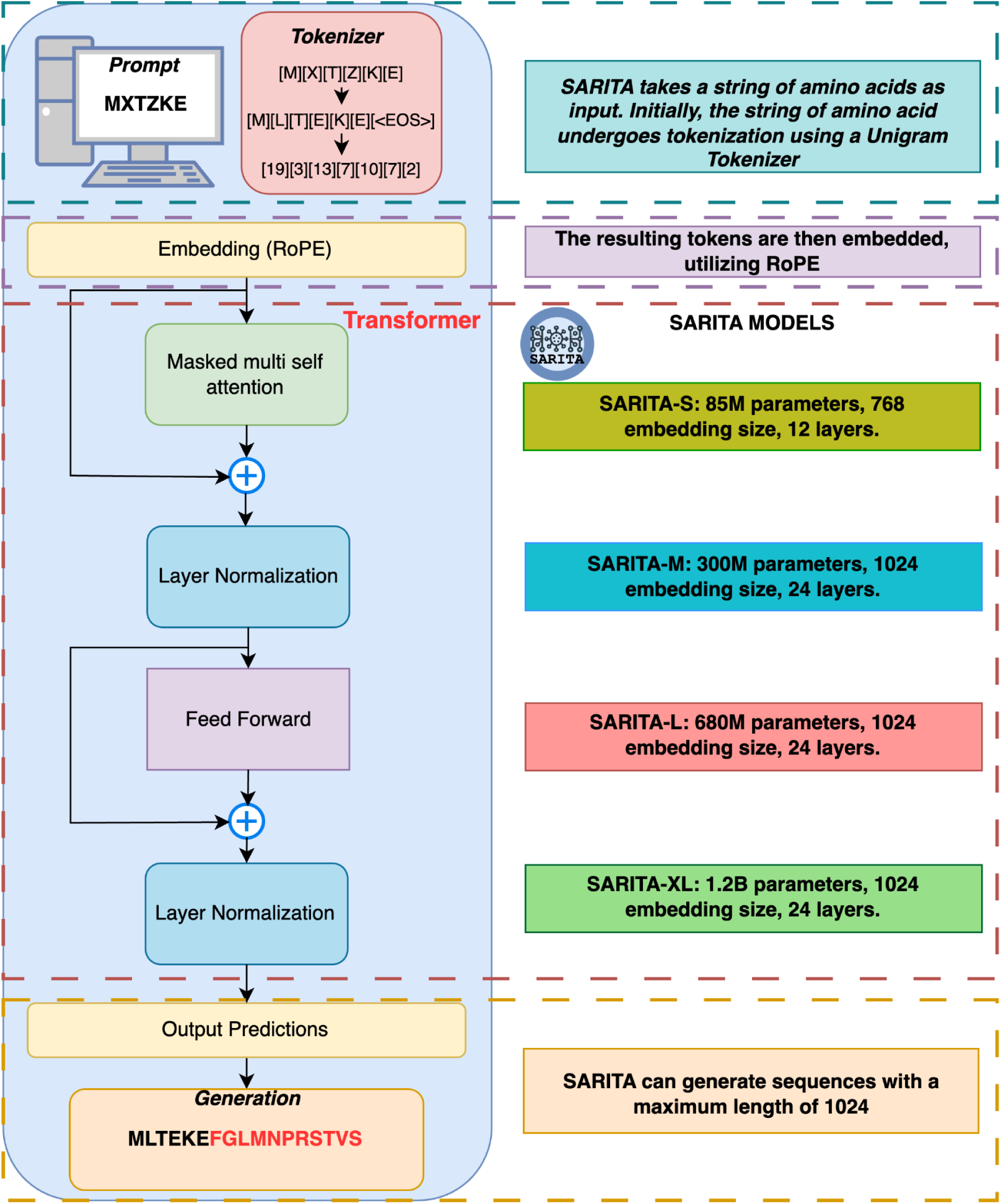
General architecture of SARITA. (Image realized with draw.io [3].)

### Training set

We downloaded 16,187,950 Spike protein sequences from the GISAID database. The sequences come from samples collected from December 19th, 2019 to November 30th, 2023. We select proteins that i) are derived from genomes marked as high coverage in GISAID i.e., genome with <1% Ns and <0.05% unique amino acid mutations (not seen in other sequences in database) and no insertion/deletion unless verified by submitter; ii) have a length of 1,273±30 amino acids, thus representing a complete Spike sequence; iii) are collected from a human host; iv) have a lineage assigned, determined by the Pango algorithm [ref.]; v) do not contain letters representing non-unique or non-standard amino acids (i.e., “X”, “B”, “Z”, ”J”, “U”, “O”); and vi) have an associated submission date complete with year, month, and day [6].

After filtering, 4,952,389 high-quality Spike sequences remain. We select sequences collected between December 19th, 2019, and February 28st, 2021 as the training set, resulting in a total of 793,732 available sequences. As the distribution of the unique sequences is far from uniform, with some unique sequences having a very high number of copies, we implement a subsampling strategy to mitigate this bias. Our rationale is that overrepresented sequences can lead SARITA to skew the generated synthetic sequences toward the overrepresented ones, limiting its ability to produce novel and previously unseen sequences [13]. To balance the dataset we use the mean number of copies of unique sequences (n=25) as a an upper threshold for the number of copies of the very same sequence, i.e., any unique sequence present with a number of copies (instances) above the initial mean value is subsampled to 25 copies. The Gini index for the unique sequence copy number decreases from 0.93 to 0.07 after resampling [17]. A total of 107,017 sequences (32,376 unique) are eventually available for training (of these, 10% for training validation).

### Test set

The unique GISAID sequences with a collection date spanning from March 2021 to November 2023 (145,059 in total) are used as a test set. Importantly, by design *only* the Alpha, Beta and Gamma variants are included in the training set, while other known variants, such as, Delta, Omicron and Kappa, are only present in the test set. We can therefore validate the ability of SARITA in predicting Variants of Concern (VOCs) like Delta and Omicron, as well as their sub-lineages or Variants of Interest (VOIs), including Kappa, Iota, and Zeta that emerged after March 1st, 2021 (**Figure S1**). This test set is used to measure the performance of SARITA (see Evaluation strategy, below).

### Continual learning

We apply continual learning [11] to fine-tune the general-domain pretrained model (RITA) with external, domain-relevant resources, enabling the model to incrementally acquire new knowledge. For this phase, we configure training with 10 epochs and a batch size of 8 for both training and validation, saving model checkpoints at the end of each epoch and evaluating the model on validation data after every epoch. The training is optimized with floating-point precision 16 (fp16) on CUDA for efficiency. We utilize University of Florida Hipergator computing resources (six A100 GPUs, 100 GB of RAM, and 10 CPUs per task).

### Competing approaches

We compare SARITA with several other approaches (Table 1). As baseline, we implement three simple random sequence generation methods: The first samples each amino acid position entirely at random (“Rand”); the second samples each position based on amino acid frequency in the training set (“Rand Fr”); and the third uses PAM30 weights for sampling (“Rand PAM30”). We also evaluate the performance of the following LLMs: SpikeGPT2, RITA-S, RITA-M, RITA-L, and RITA-XL [13]. SpikeGPT2 is an autoregressive model with 738 million parameters derived from ProtGPT2 [18] initially trained on the UniRef50 dataset to learn general protein structures, fine-tuned on 20,000 SARS-CoV-2 Spike protein sequences, specifically focusing on sequences gathered prior to May 2021. The fine-tuning enables SpikeGPT2 to adapt to SARS-CoV-2 mutation patterns, with modifications applied to select layers only, preserving core protein knowledge while specializing in SARS-CoV-2 mutations. RITA is a group of four autoregressive generative models for protein sequences, scaling up to 1.2 billion parameters, trained on more than 280 million protein sequences from UniRef100 [19], MGnify [20], and Metaclust [21].

**Table 1:** Comparison between SARITA and the competing models, in terms of i) number of model parameters, ii) computational resources used for prediction, and iii) number (percentage) of high-quality generated sequences.

| Model | # of parameters | Computational resources | # of evaluated sequences (after filtering) |
| --- | --- | --- | --- |
| Rand | - | 1 Core, 1.41MB | 11,300 (100%) |
| Rand FR | - | 1 Core, 1.41MB | 11,300 (100%) |
| Rand PAM30 | - | 1 Core, 243MB | 11,300 (100%) |
| RITA S | 85 million | 1 A100, 1 Core, 6.06 GB | 11,300 (100%) |
| RITA M | 120 million | 1 A100, 1 Core, 6.20 GB | 11,300 (100%) |
| RITA L | 300 million | 1 A100, 1 Core, 6.07 GB | 11,300 (100%) |
| RITA XL | 1.2 billion | 1 A100, 1 Core, 6.05 GB | 11,300 (100%) |
| SpikeGPT2 | 738 million | 1 A100, 1 Core, 6.07 GB | 339 (3%) |
| SARITA S | 85 million | 1 A100, 1 Core, 6.07 GB | 11,074 (98%) |
| SARITA M | 300 million | 1 A100, 1 Core, | 11,074 (98%) |
|  |  | 6.22 GB |  |
| SARITA L | 600 million | 1 A100,1 Core,<br>6.05 GB | 10,961 (97%) |
| SARITA XL | 1.2 billion | 1 A100,1 Core,<br>6.05 GB | 11,187 (99%) |

### Evaluation strategy

The evaluation strategy involves assessing the performance of various models in generating S1 Spike proteins. To generate the sequences to be tested (**Figures 2 and 3**), we used all the 113 unique prompts of 14 amino acids found in the training set preceding the S1 Subunit. For each prompt, we use the SARITA, SpikeGPT2, RITA, and random models to generate 100 sequences, each with a length of 686 amino acids, including the prompt, thus matching the S1 length, spanning from position 15 to 686 [1][2].

**Figure 3:**
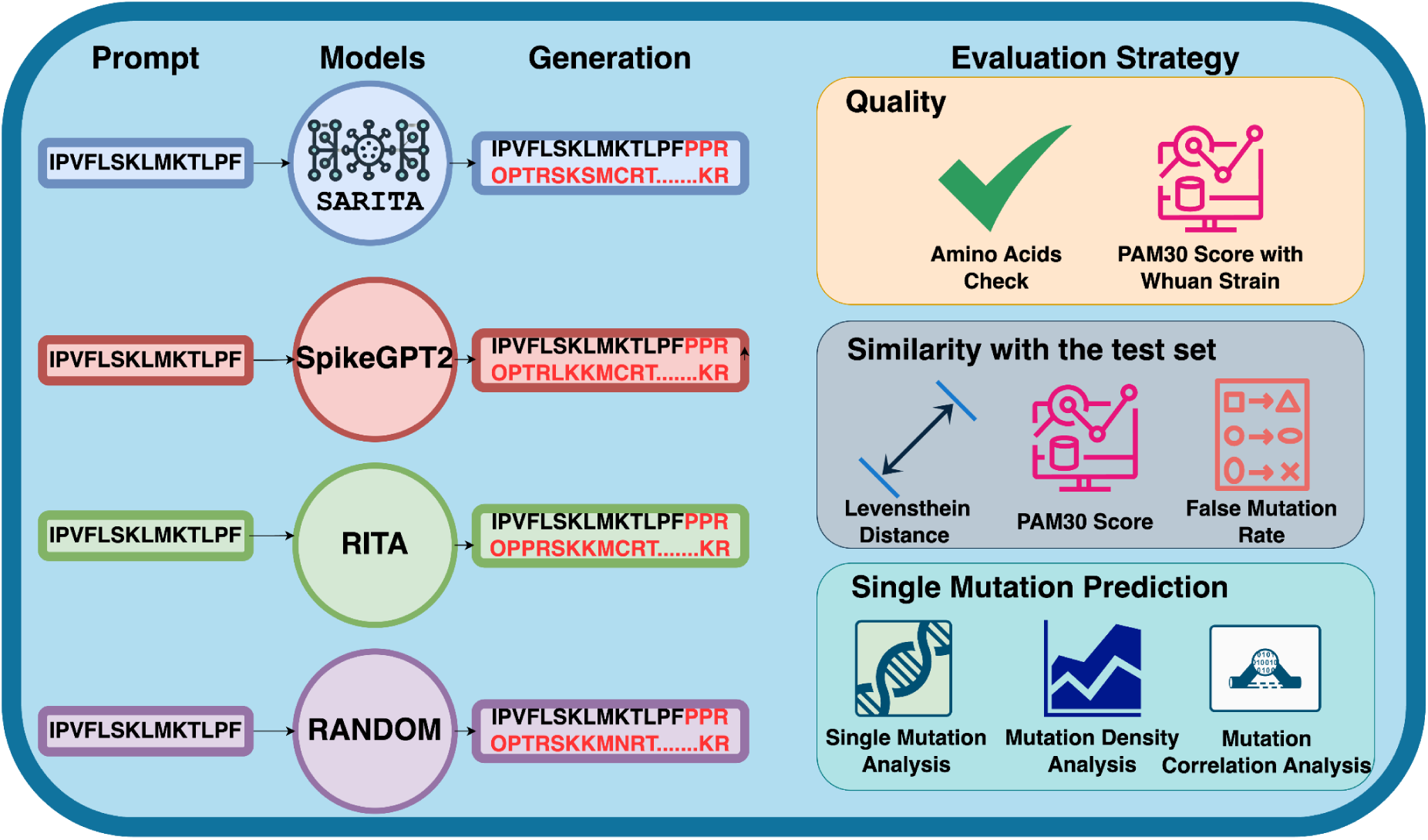
Evaluation Strategy: Schematic of the validation strategy. The 14-amino-acid prompt is used to generate S1 subunit sequences of 686 amino acids. The 113 unique prompts available in the training set are used to generate 11300 sequences per model. The evaluation covers three key aspects: <u>Quality</u>, checking for presence of non-unique or non-standard amino acid; and similarity with the Wuhan strain (PAM30 score); <u>Similarity</u>, assessed comparing the generated sequences to the test set with Levenshtein distance, PAM30 score, and false mutation rate; and Single Mutation Prediction, evaluated through single mutation analysis, mutation density analysis, and correlation mutation analysis on the test set. (Image realized with draw.io [3].)

To evaluate the performance of SARITA and its competing approaches, we assess three main aspects: i) sequence quality, i.e., are the generated sequences composed of valid amino acids? ii) similarity, i.e., are the generated sequences similar to the ones in the test set? And iii) single-point mutation prediction, i.e., are the generated mutations biologically plausible?

### Quality of generated proteins

The quality of the generated sequences is checked by the presence of invalid amino acids, represented by characters not corresponding to the twenty canonical amino acids (i.e., “X”, “B”, “Z”, ”J”, “U”, “O”). We further examine the quality of the synthetic sequences by measuring their similarity to the original Wuhan strain. Our rationale is that higher dissimilarity implies less realism. To measure similarity, we use alignment (Needleman–Wunsch, Biopython package version 1.80, default parameters) via the PAM30 scoring matrix. PAM30 assesses evolutionary similarity by assigning scores to amino acid substitutions based on their likelihood over short evolutionary distances and is optimized for closely related protein sequences. Higher PAM30 scores indicate stronger similarity to the Wuhan strain, suggesting conservation of structural or functional properties, while lower scores imply greater divergence, potentially highlighting adaptive mutations or functional shifts.

### Sequence similarity with the test set

To assess if the approaches are capable of modeling viral evolution, i.e., to predict future sequences, we measure the similarity between the generated sequences and test set. We compute the Levenshtein distance (LD) [22], which is commonly used to compare strings, representing the minimal cost of transforming one string into another through deletion, insertion, or substitution . For each generated sequence, we calculate LD against all the test sequences, recording both the minimum and median distance. In the case of SpikeGPT2, we use the full generated protein length to calculate LD.

We further assess viral evolution prediction via PAM30 scoring as described above: We compare each generated sequence with the test set, and retain the best score. Similarity if further investigated by calculating the False Mutation Rate (FMR), i.e., the presence of mutations in the synthetic sequences that are not matched by the best alignment in the test set. Given a synthetic sequence *S* and its best matching test set sequence *M*, we calculate the FMR as

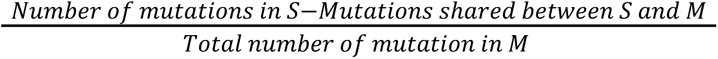

FMR is the ratio between the number of mutations found in *S*, but not present in *M*, divided by the total number of mutations in *M* (with *S* and *M* mutations assessed via alignment with the original Wuhan S1).

### Single-point mutation prediction

To evaluate single-point mutation prediction, we examine whether the mutations of the synthetic sequences show a positional correspondence with the ones of the training and the test sets. We compute the density difference between mutations in synthetic and real sequences. For each S1 position, we examine how mutations are generated by SARITA and estimate the density using kernel smoothing, a non-parametric method producing a continuous curve from discrete data points without imposing strict assumptions on the underlying distribution shape [7]. We set an adjustment parameter of 0.1 to control the kernel width. Lower values create a more detailed but less smooth curve, while higher values produce a smoother, less detailed curve. To adjust for observation frequency, we use the base-10 logarithm of the mutation frequency, which reduces the impact of extremely high frequency values and enhances interpretability. This comprehensive approach provides insights into the distribution and biological relevance of synthetic mutations compared to real mutation patterns. We calculate the population frequency of S1 mutations for each position in the sequences generated by SARITA and those in the training and test sets. Then, we computed Pearson’s correlations.

### Availability

Both training and test set GISAID sequence IDs, along with the four final models trained on the whole set (training + test, up to August 2024) are available on Hugging Face: https://huggingface.co/SimoRancati

## Results

SARITA has been trained on 612,759 SARS-Cov-2 complete, high-quality Spike protein sequences uploaded to the GISAID database [14]. SARITA takes as an input the first 14 amino acids of the Spike S1 subunit, and generates a synthetic S1 subunit sequence (**Figure 3**). To estimate its performances, we split GISAID data into a training set (December 19th 2019 to February 28th 2021) and test set (March 1st 2021 to November 8th 2023). Information about training times and batches are reported in the supplementary in **Table S2**.

All the 113 unique 14-amino-acid prompts present in the training set are used to generate 11,300 new sequences, which were then compared to the real-world test set. We evaluate the generated synthetic sequences across three main objectives: i) Quality; ii) sequence similarity with the test set; and iii) single-point mutation prediction. Quality is assessed by checking for the presence of undefined or non-standard amino acids and biological alignment against the original Wuhan S1. Sequence similarity with the test is evaluated with LD, PAM30 scores, and FMR. Single-point mutation prediction is evaluated by examining the ability to replicate position-dependent mutational patterns.

### Quality: SARITA generates high-quality, synthetic SARS-Cov-2 Spike S1 sequences

For each of the 113 possible prompts of the training set we generate 100 synthetic sequences for a total of 11,300 sequences for each SARITA model (S, M, L, XL). We compared the results against an equivalent number of S1 sequences generated by competing approaches: two other LLMs (SpikeGPT2 and RITA); and three baseline approaches (“Rand”, “Rand Fr”, “Rand PAM30”).

If a generated sequence shows one or more uncharacterized amino acids, it is not considered high-quality [1,4]. Another measure of quality is related to the sequence length: We consider a sequence with a length of 686±10 to be high-quality. SARITA generates high-quality sequences at least 97% of the times (**Table 1, Figure S2**). RITA models provide a better performance, with 100% high-quality generated sequences. Instead, only 3% of SpikeGPT2-generated sequences (339 out of 11,300) are high-quality. Additionally, none of the sequences generated by SpikeGPT2 actually match the expected S1 length, despite the fact that it was instructed to match the length of the S1 subunit. Specifically, the median length of SpikeGPT2-generated sequences is 2,003 amino acids, approximately 192% longer than the expected S1 subunit length (**Figure S2**).

SARITA outperforms competing methods in generating sequences closely related to the original Wuhan strain, therefore indicating higher biological plausibility and realism, as shown by the median PAM30 score (**Figure 4**). Specifically, SARITA produces synthetic sequences with median PAM30 scores ranging from 5,191 (SARITA S) to 5,278 (SARITA M), indicating high similarity to the original sequence. In contrast, all competing methods generate sequences with significantly lower PAM30 scores, indicating a lower-quality alignment with biological S1 sequences. Their median scores range from 661 (Rand PAM30) to 2,062 (SpikeGPT2).

**Figure 4.**
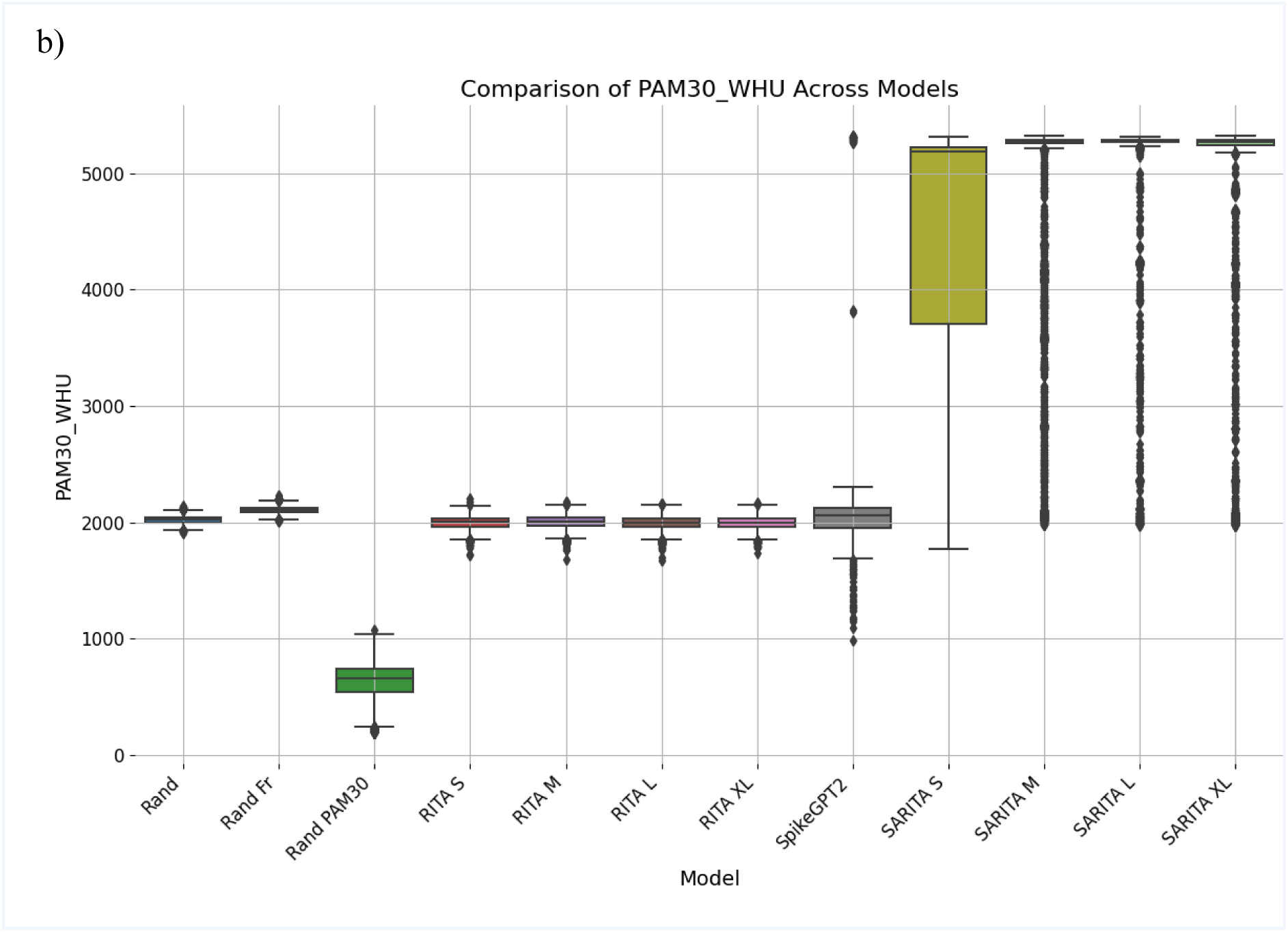
Boxplot showing the PAM30 distribution scores computed on the synthetic sequences (higher PAM30 score means that the sequences are strongly related)

### Sequence similarity with the test set: SARITA outperforms competing approaches in matching viral evolution

We assess if the models produce plausible future S1 sequences, matching the ones of the test set. Taken together, low LDs suggest the synthetic sequences closely resemble the test set, indicating the model correctly predicts viral evolution. **Table 2** shows how SARITA greatly outperforms other models. RITA models, along with the random baseline models, never achieve a best LD<500, and only 1.8% of SpikeGPT2 achieve a best LD <500. In contrast, SARITA S, M, X, and XL reach a best LD<10 with 93%, 98%, 98%, and 99% of the sequences respectively. Further details about LD results are shown in **Figure S3**.

**Table 2.** Percentage of generated sequences with LD < 4, 10, 100, 500, 700, 1500, and 2000 against the best aligned test sequence.

| Model | LD<4 | LD<10 | LD<100 | LD<500 | LD<700 | LD<1500 | LD<2000 |
| --- | --- | --- | --- | --- | --- | --- | --- |
| Rand | 0% | 0% | 0% | 0% | 100% | 100% | 100% |
| Rand Fr | 0% | 0% | 0% | 0% | 100% | 100% | 100% |
| Rand PAM30 | 0% | 0% | 0% | 0% | 100% | 100% | 100% |
| RITA S | 0% | 0% | 0% | 0% | 100% | 100% | 100% |
| RITA M | 0% | 0% | 0% | 0% | 100% | 100% | 100% |
| RITA L | 0% | 0% | 0% | 0% | 100% | 100% | 100% |
| RITA XL | 0% | 0% | 0% | 0% | 100% | 100% | 100% |
| SpikeGPT2 | 0% | 0% | 0% | 1.8% | 5.6% | 75% | 100% |
| SARITA S | 31.7% | 93.2% | 99.3% | 100% | 100% | 100% | 100% |
| SARITA M | 86.% | 97.9% | 98.9% | 100% | 100% | 100% | 100% |
| SARITA L | 63.5% | 98.4% | 98.4% | 100% | 100% | 100% | 100% |
| SARITA XL | 53.7% | 98.6% | 98.7% | 100% | 100% | 100% | 100% |

Best SARITA median PAM30 scores range from 5,210 (SARITA S) to 5,302 (SARITA M), indicating a high similarity with the sequences in the test set. In contrast, the competing model scores range from 670 (Rand PAM30) to 2,102 (SpikeGPT2).

We also compute the FMR, or the frequency at which SARITA introduces mutations in synthetic sequences that are not corresponding with the best alignment with test set sequences. The results (**Figure 5**) indicate that SARITA outperforms existing methods in producing realistic synthetic sequences, achieving a minimum of 0.5 (median) with SARITA L, whereas the best result among all competing methods is 0.98 (median). Additionally, while other methods introduce hundreds of mutations in a single sequence (SpikeGPT2 achieves the best median with 438 mutations), making it highly unrealistic, SARITA maintains a controlled mutation count, with the median ranging from 38 for SARITA S, to just 14 for SARITA L.

**Figure 5:**
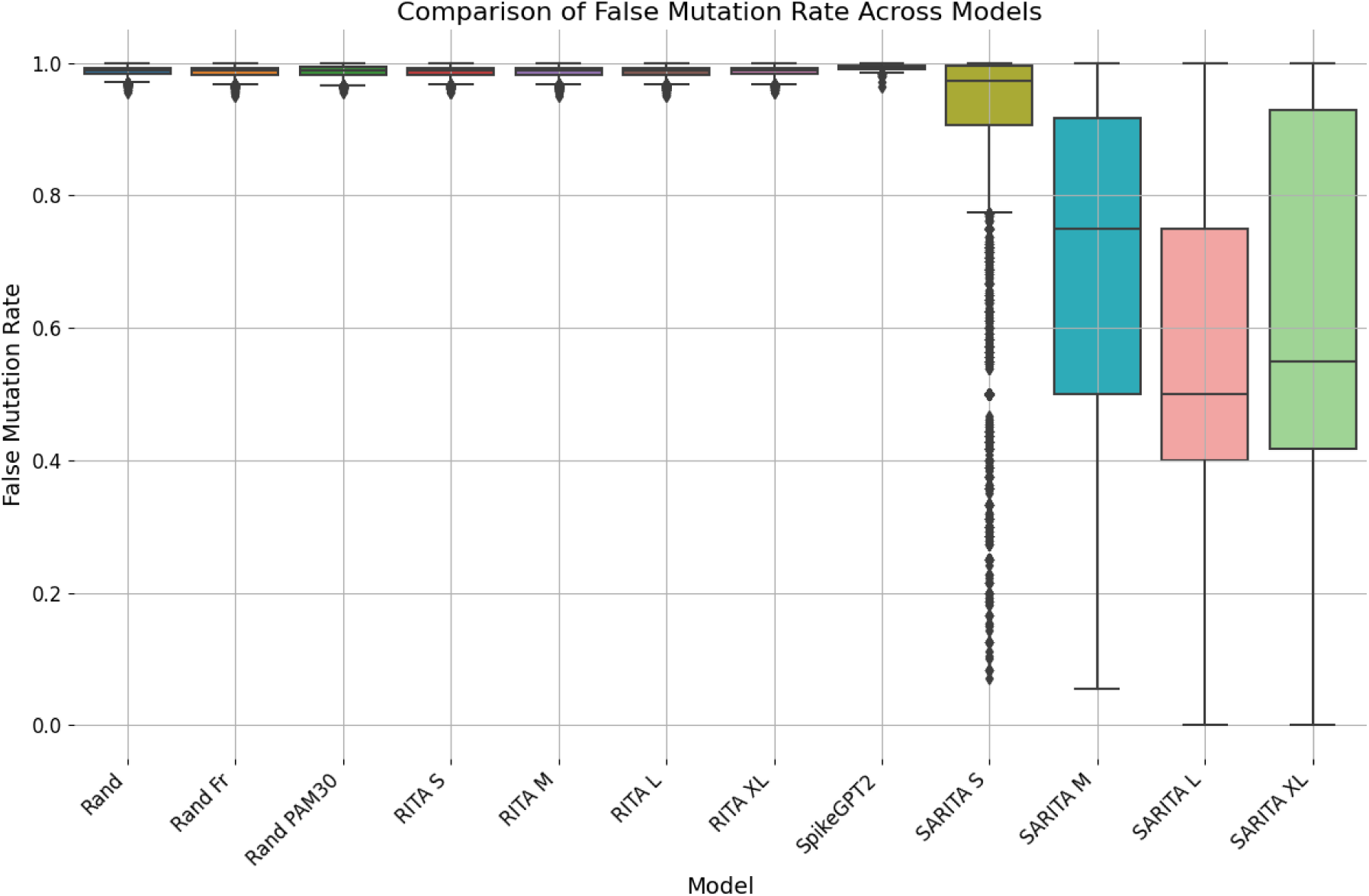
Boxplot displaying the distribution of mutation false mutation rate (FMR) across synthetic sequences. The FMR is calculated as the proportion of additional mutations—those not present in the best-aligned sequence—relative to the total mutations generated by the model. SARITA shows superior performance over the comparison model.

### Single-point mutation prediction: SARITA correctly reproduces S1 mutation evolution

To evaluate the ability to generate realistic VOCs such as Delta and Omicron, and VOIs, like Iota and Zeta, we analyze the generated mutations, comparing them to the key VOC and VOI mutation. SARITA successfully reproduce key mutations associated with different variants, including L212I, an important mutation in RBD characterizing Omicron variants [23]; R158L, a mutation altering the secondary structure of the protein, characteristic of Iota [24]; and T19L, a common mutation in Delta variants and Omicron sublineages such as BA.2 [25]. Additionally, SARITA generated T95P, a mutation seen in both Delta and Iota [26]. Another mutation identified by SARITA is E406K, characterizing Delta variants and abrogating neutralization mediated by the REGEN-CoV therapeutic monoclonal antibody (mAb) COVID-19 cocktail and the cilgavimab (AZD1061) mAb [27]. Remarkably, SARITA can generate sequences with mutations from newer VOCs, evolved since March 2021, which are not present in the training set. These include Delta, Omicron, Epsilon, Eta, Iota, and Kappa, highlighting the capability of SARITA to simulate mutations in emerging variants.

**Figure 6** shows distinct patterns in mutation generation across the SARITA models. Notably, the proportion of novel mutations, representing mutations not present in either the training or test data, increases with model size, rising from 48% in SARITA-S to 55% in SARITA-M, and stabilizing at 54% for SARITA-L and SARITA-XL. This trend suggests that larger models are better equipped to generate novel mutations, an essential attribute for enhancing generalization capabilities. To assess the plausibility and realism of novel mutations, we calculate their PAM30 scores and compare the score distribution with those from the training and test sets (**Figure 7**). The figure shows how the PAM30 score of the generated mutations is very close—with a median difference of only 1 point—to the mutations in the training and test sets.

**Figure 6:**
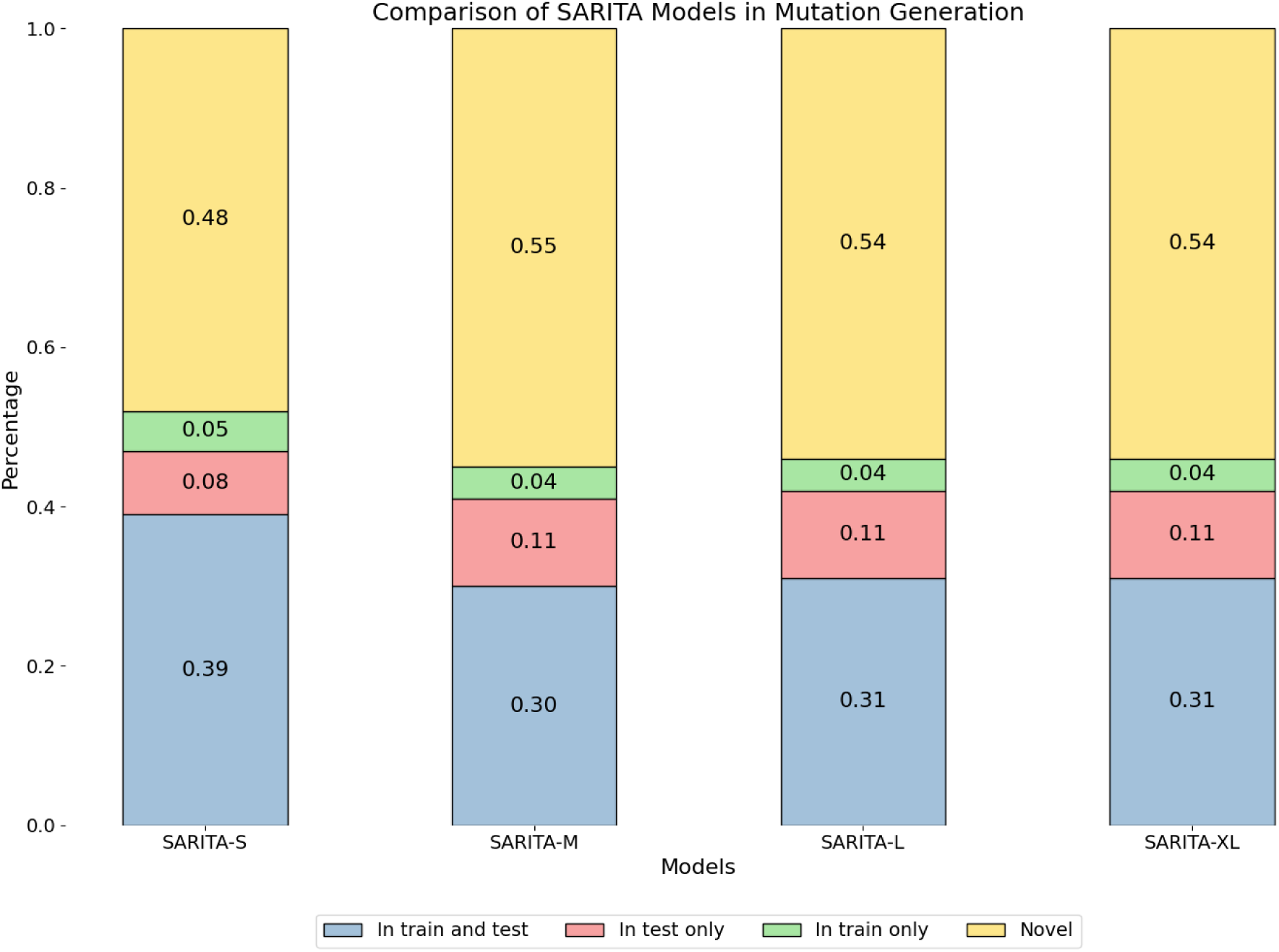
Comparison of SARITA Models in Mutation Generation. The stacked bar chart represents the distribution of mutations across four SARITA models Each bar is divided into four categories: present in both training and test sets (blue); present in test set only (red); present in training set only (green); and novel, i.e., not present in the training nor the test sets (yellow).

**Figure 7:**
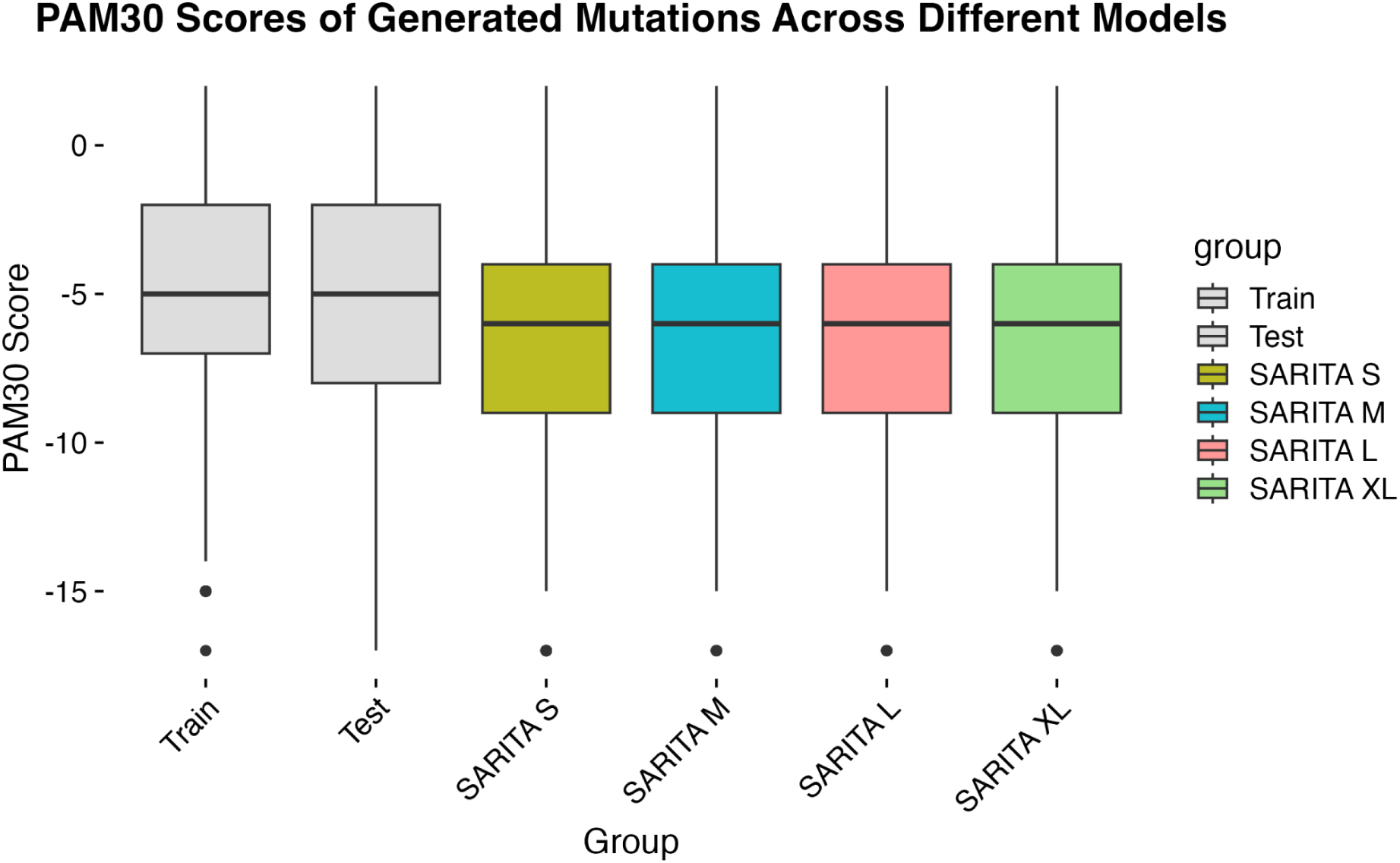
Boxplot displaying the distribution of PAM30 score across synthetic mutations sequences. The PAM30 score is calculated through the alignment with the S1 Wuhan Strains.

Lastly, we calculate both the frequency and relative density of mutation changes at each position in sequences generated by SARITA and those in the training and test sets. **Figure 8** shows SARITA M predictions closely align with the observed distribution of variants across positions in the SARS-CoV-2 S1 subunit, capturing both pre- (training) and post-February 2021 (test) distributions. Additionally, we calculate the Mean Square Error (MSE) between the test set variant density and the SARITA-generated variant density, obtaining -8.94e-08, -6.57e-08, -7.22e-08, and -8.46e-08 for S, M, L, and XL models respectively. These results indicate that SARITA M offers the closest approximation to the test set distribution among the model sizes. **Figure 9B** further shows the strong positive correlation (0.7, p < 2.2e-16, Pearson) between the mutation frequencies in sequences generated by SARITA M (Y-axis) and those of the test set (X-axis). Correlation results for SARITA of other sizes (S, L, XL) are detailed in **Table S3**.

**Figure 8:**
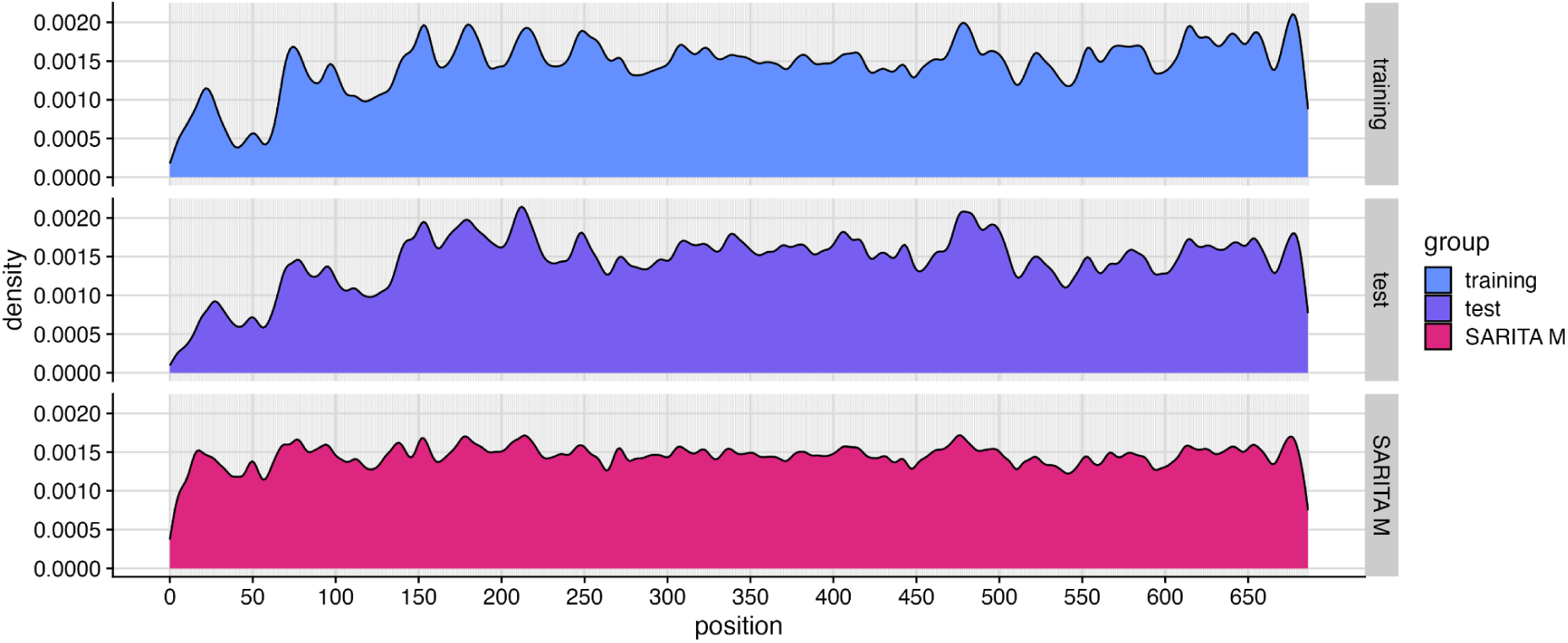
Gaussian kernel density estimate plots show that the observed distribution of variants across positions in the SARS-CoV-2 S1 Subunit both pre- (training) and post-February 2021 (test) are well approximated by SARITA M.

**Figure 9:**
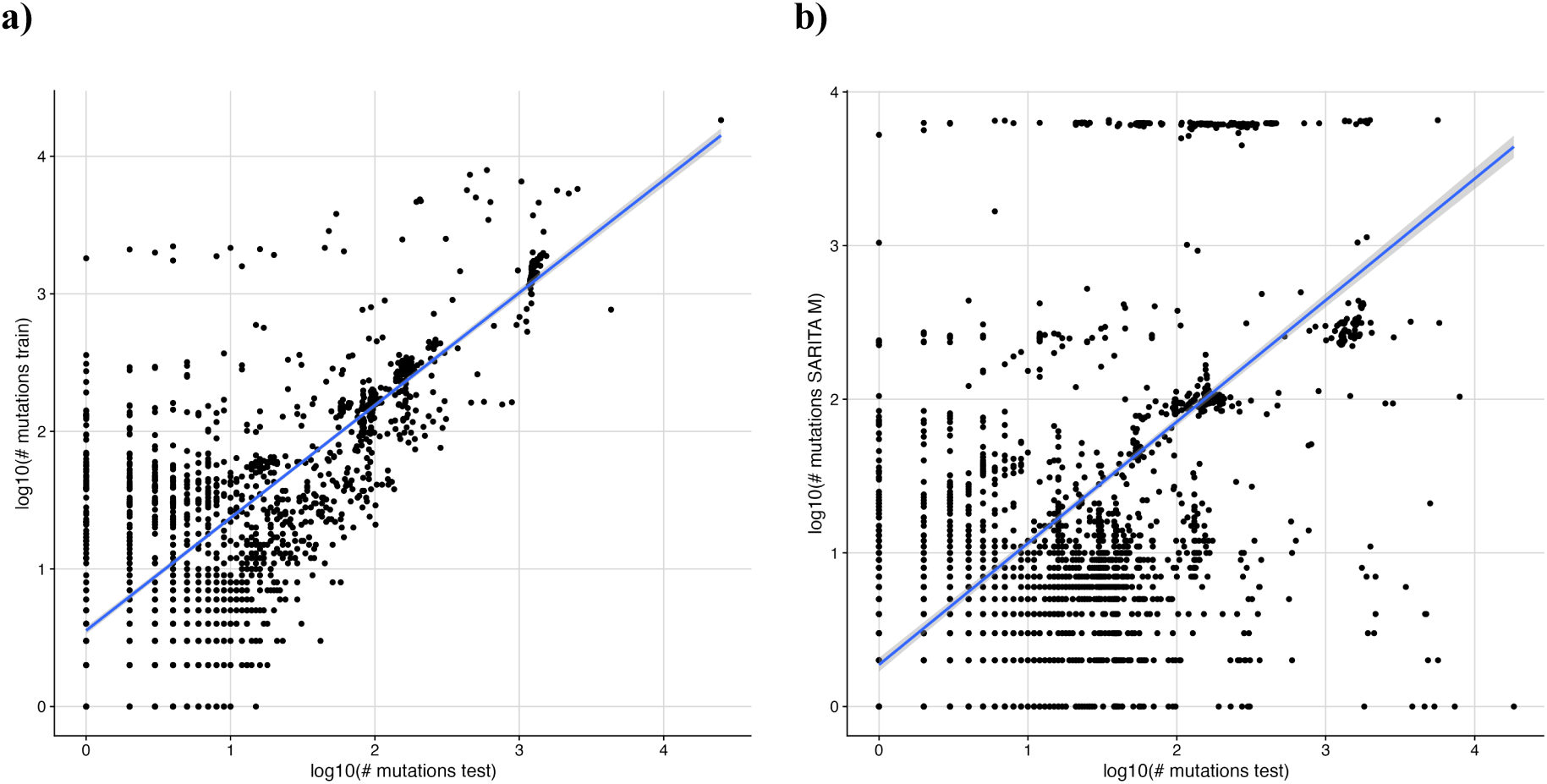
Correlation of S1 subunit mutations. **a)** Correlation of mutation frequency in logarithmic scale between training (x-axis) and test (y-axis) set. **b)** Correlation of mutation frequency in logarithmic scale between SARITA-M sequences (x-axis) and test (y-axis) set.

## Discussion

The results confirm SARITA as a highly specialized LLM for generating biologically meaningful synthetic SARS-CoV-2 S1 sequences. SARS-CoV-2 has continued to mutate and evolve since the pandemic onset, challenging public health responses and emphasizing the importance of predictive models that can *anticipate* viral changes. By generating synthetic sequences that align closely with future real-world mutational patterns, SARITA addresses a critical need for proactive measures in viral genomics, providing a tool that can predict natural viral evolution in SARS-CoV-2 Spike protein.

One of the most notable outcomes is the ability to reproduce both known and future mutational patterns among VOCs and VOIs. For example, SARITA accurately reproduces mutations such as L212I, R158L, and T19L, associated with specific variants like Omicron and Delta[23], [24], [25]. A particularly valuable aspect of SARITA is its parsimonious and realistic mutation generation: Unlike other generative models, SARITA maintains a low FMR and does not introduce an excessive number of mutations within a single synthetic S1. This reasonable mutational load, similar to the one of real-world data, is crucial for creating synthetic sequences that are biologically plausible. By maintaining this balance, SARITA supports studies aimed at understanding the evolutionary trajectory of SARS-CoV-2 without sacrificing biological integrity, making it a more reliable model for predicting variant emergence.

In terms of model comparison, SARITA outperforms (**Figure 10**) alternative generative models, including SpikeGPT2 and RITA, by generating sequences with lower LD and higher PAM30 scores relative to real-world viral sequences. One of the key innovations is its generation of novel mutations that are not present in either the training or test sets, yet are highly realistic, reflecting the ability to explore mutational landscapes beyond known variants. The fact that larger SARITA models, such as SARITA-L and SARITA-XL, generate a higher proportion of novel mutations compared to smaller models supports the idea that scaling model parameters enhances generative diversity. This feature is promising for applications where anticipating novel mutations is necessary, such as in the early identification of potentially dangerous variants that could arise due to selective pressures or recombination events in viral genomes.

**Figure 10:**
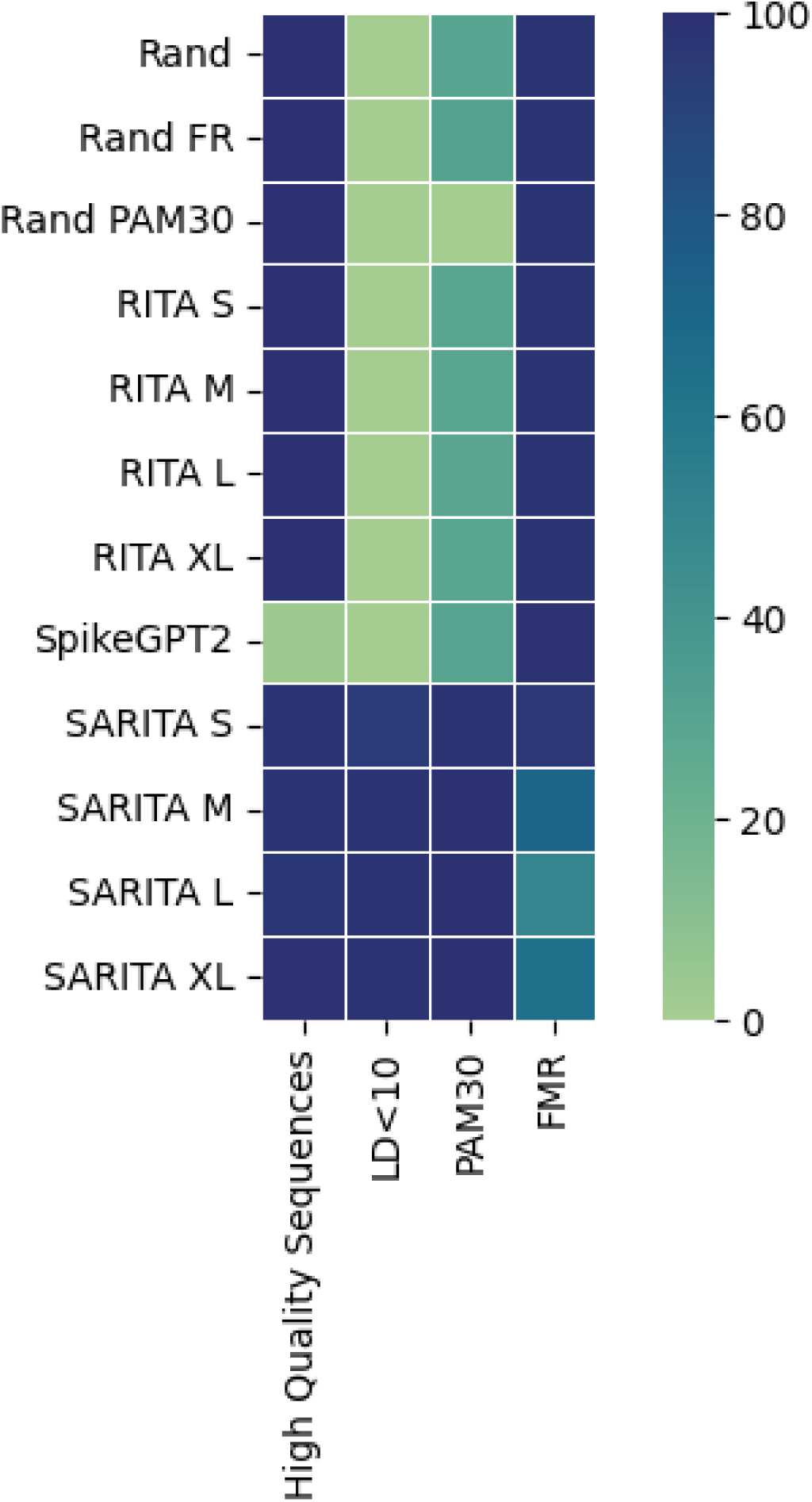
Heatmap illustrating the false mutation rate (FMR) across various models and configurations, including random baseline models (Rand, Rand FR, Rand PAM30), RITA models (S, M, L, XL), SpikeGPT2, and SARITA models (S, M, L, XL). Each model is evaluated based on different metrics: Percentage of high-quality sequences, percentage of sequences with LD<10 (LD<10), normalized value of PAM30 with the original Wuhan strain (PAM30), and FMR.

**Figure 10** shows that the worst method for generating synthetic S1 subunits is Rand_PAM30. Rand_PAM30 might underperform the other random models due to overfitting specific amino acid substitutions favored by PAM30, which are too general and do not very well fit the SARS-CoV-2 genome. PAM30 introduces biases toward particular substitutions, potentially limiting its general applicability. In contrast, the basic Rand model, being fully random, avoids these biases and achieves relatively better results.

Despite its strengths, SARITA has limitations that should be addressed in future studies. While the model shows high efficacy in generating SARS-CoV-2 sequences, adapting SARITA to other viruses with different mutation dynamics and selection pressures would require substantial re-training and refinement. Additionally, SARITA predictions are limited by the quality and completeness of the data in GISAID, meaning that any biases or gaps in this data source could influence the output.

Future research could expand on SARITA by exploring its adaptability to other coronaviruses or respiratory pathogens. Integrating SARITA outputs into a broader predictive framework that includes epidemiological data, host-pathogen interaction studies, and environmental factors could provide a holistic view of viral evolution, supporting comprehensive and adaptive public health strategies.

## Conclusions

We present SARITA, a generative LLM tailored on SARS-CoV-2 Spike S1. SARITA outperforms other models in generating high-quality, realistic sequences, and shows the capability of correctly anticipating viral evolution. SARITA could contribute to a more proactive framework in genomic surveillance, equipping researchers with a tool to anticipate and prepare for future SARS-CoV-2 variants.

## Acknowledgements

This study has been funded in part by the NIH grant NIAID R01 AI170187; and by the Artificial Intelligence and Complex Computational Research Award, University of Florida.

