## Supplementary for "SARITA: A Large Language Model for Generating the S1 Subunit of the SARS-CoV-2 Spike Protein"

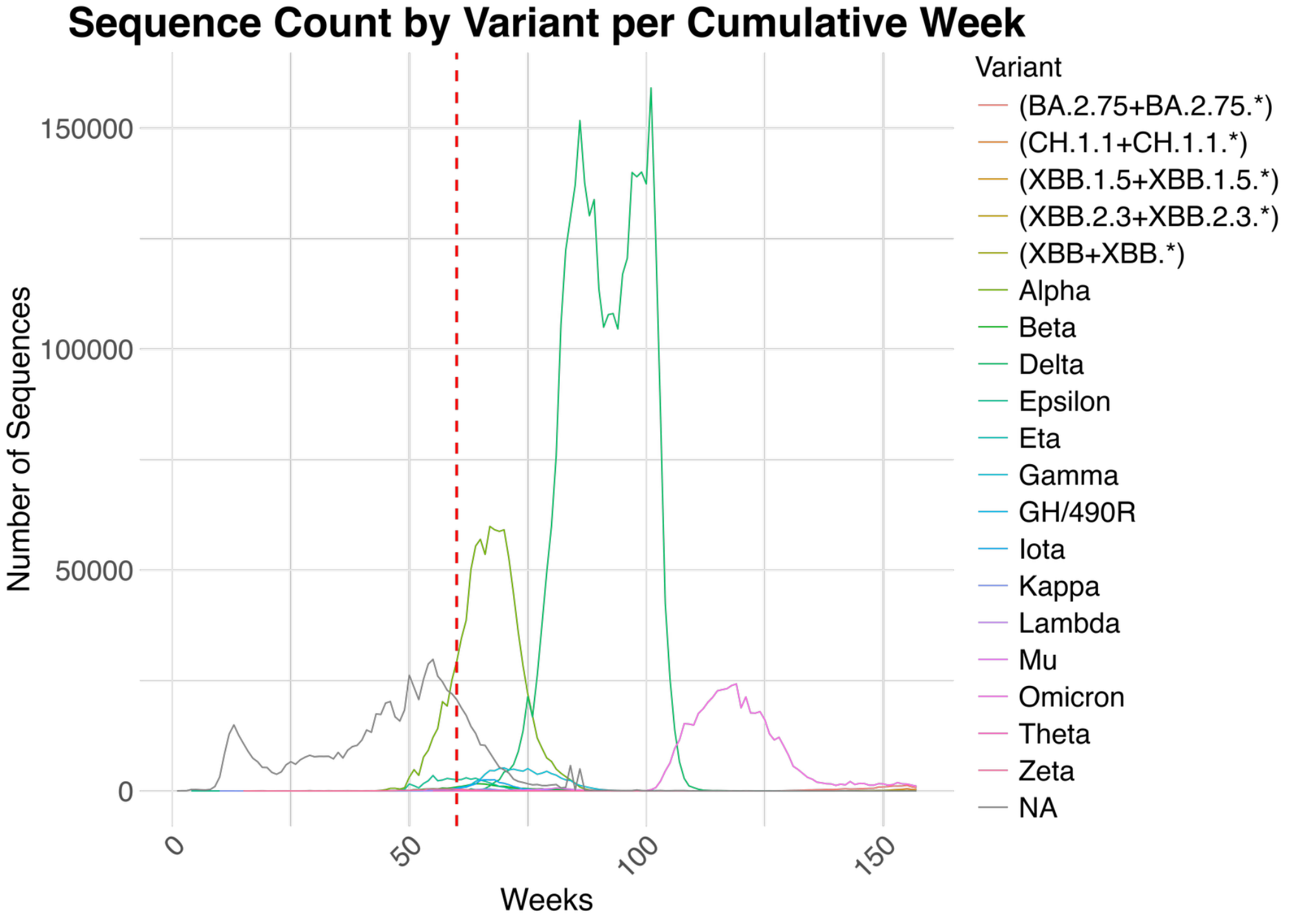

**Figure S1:** The number of sequences uploaded to GISAID from December 2019 to November 2023 for relevant VOCs/VOIs. The red line marks the division between the training set (left) and the test set (right).

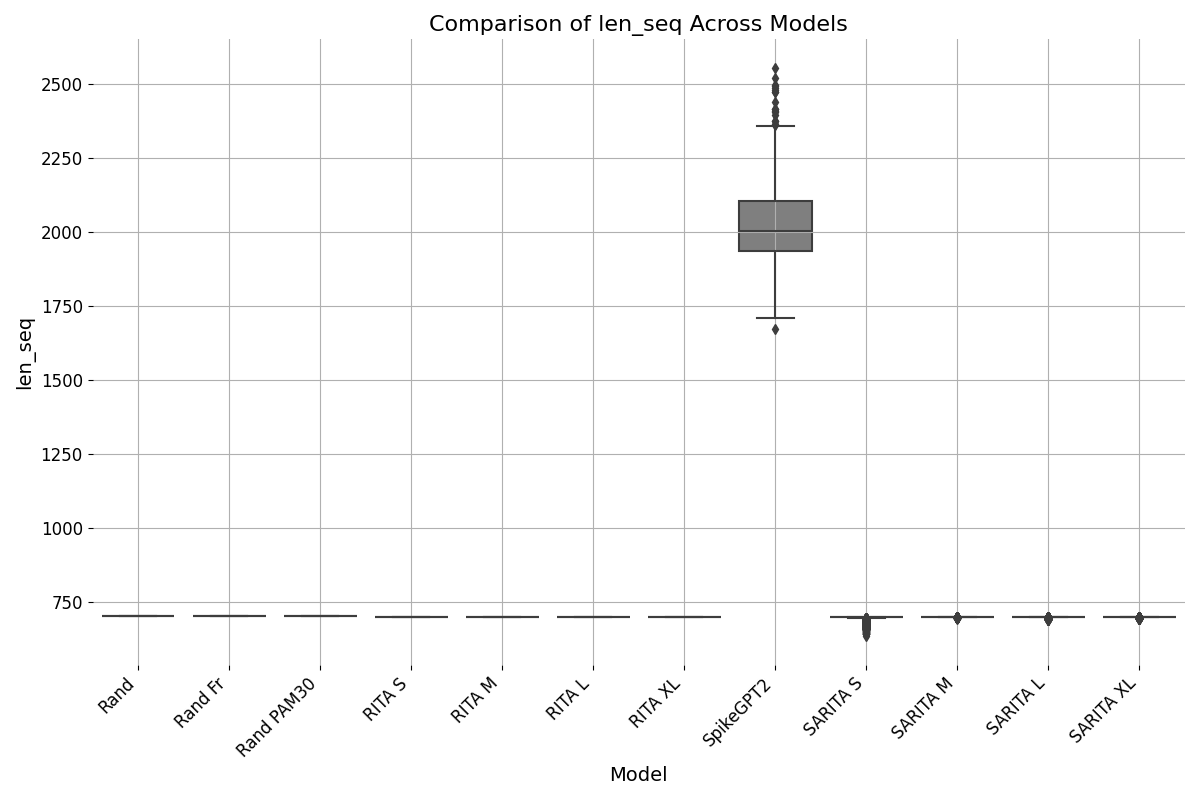

**Figure S2:** Boxplot illustrating the distribution of sequence lengths (len_seq) generated in synthetic sequences. Notably, SpikeGPT2 does not adhere to user-defined limits for creating sequences with a fixed length.

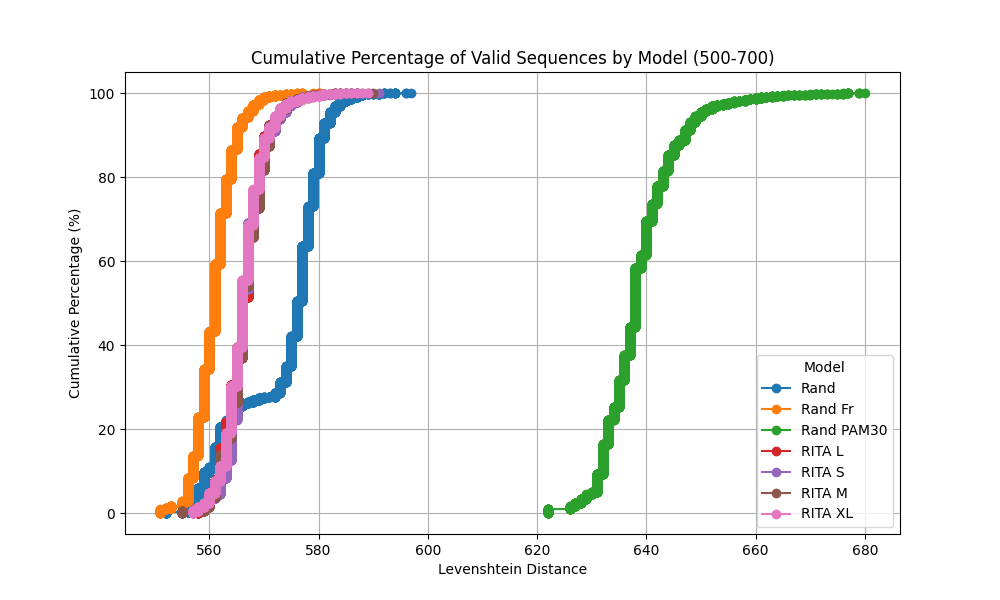

**Figure S3**: Cumulative percentage of sequences as a function of Levenshtein distance (500-700 range) for random methods (Rand, Rand Fr, Rand PAM30) and RITA models (RITA S, RITA M, RITA L, RITA XL).

| **Token Content** | **Token ID** |
| --- | --- |
| <PAD> | 1 |
| <EOS> | 2 |
| L | 3 |
| A | 4 |
| G | 5 |
| V | 6 |
| E | 7 |
| S | 8 |
| I | 9 |
| K | 10 |
| R | 11 |
| D | 12 |
| T | 13 |
| P | 14 |
| N | 15 |
| Q | 16 |
| F | 17 |
| Y | 18 |
| M | 19 |
| H | 20 |
| C | 21 |
| W | 22 |
| B | 23 |
| O | 24 |
| U | 25 |
| Z | 26 |

**Table S1**: Lookup table mapping each token content to its corresponding token ID.

|  | Training times | Batches |
| --- | --- | --- |
| SARITA-S | 6 hours, 21 minutes and 15 seconds | 8 (Training Data)  8 (Validation Data) |
| SARITA-M | 15 hours, 30 minutes, 54 seconds | 8 (Training Data)  8 (Validation Data) |
| SARITA-L | 1 day, 1 hour, 17 minutes, 45 seconds | 8 (Training Data)  8 (Validation Data) |
| SARITA-XL | 1 day, 12 hours, 32 minutes, 27 seconds | 8 (Training Data)  8 (Validation Data) |

**Table S2**: Information about training times and batches.

| **Models** | **Test - Synthetic** |
| --- | --- |
| SARITA-S | 0.66 (p value 2.2 e-16) |
| SARITA-L | 0.64 (p value 2.2 e-16) |
| SARITA-XL | 0.60 (p value 2.2 e-16) |

**Table S3**: Correlation (Spearman) of mutation frequency in logarithmic scale between SARITA sequences and test set with respective p-value.
